# Life history tradeoffs, division of labor and evolutionary transitions in individuality

**DOI:** 10.1101/2020.10.23.352948

**Authors:** Guilhem Doulcier, Katrin Hammerschmidt, Pierrick Bourrat

## Abstract

Reproductive division of labor has been proposed to play a key role for evolutionary transitions in individuality (ETIs). This chapter provides a guide to a theoretical model that addresses the role of a tradeoff between life-history traits in selecting for a reproductive division of labor during the transition from unicellular to multicellular organisms. In particular, it focuses on the five key assumptions of the model, namely (1) fitness is viability times fecundity; (2) collective traits are linear functions of their cellular counterparts; (3) there is a tradeoff between cell viability and fecundity; (4) cell contribution to the collective is optimal; and (5) there is an initial reproductive cost in large collectives. Thereafter the chapter contrasts two interpretations of the model in the context of ETIs. Originally, the model was interpreted as showing that during the transition to multicellularity the fitness of the lower-level (the cells) is “transferred” to the higher level (the collective). Despite its apparent intuitiveness, fitness transfer may obscure actual mechanisms in metaphorical language. Thus, an alternative and more conservative interpretation of the model that focuses on cell traits and the evolutionary constraints that link them is advocated. In addition, it allows for pursuing subsequent questions, such as the evolution of development.

## 1. Introduction

Division of labor is an old concept. One can find the basic idea in the Republic of Plato:

> Things are produced more plentifully and easily and of better quality when one man does one thing which is natural to him and does it in the right way, and leaves other things.

Ever since Plato, a number of theorists have proposed variations on this theme with different degrees of sophistication. In The Wealth of Nations, Adam Smith (1776) tells us that ten men in a pin factory can produce about 48000 pins in a single day when he estimated that they would only produce less than 20 each, and perhaps none if ten untrained men were doing all the 18 necessary steps to produce a pin on their own. This is because each man is *specialized* in one or two steps of the pin-producing process and thus does the steps more efficiently and without the need to switch between tasks than a man performing all the steps sequentially. Without this difference in efficiency and task-switching, there would be no advantage for a man to become a specialist because it only suffices that a single of the 18 ‘types’ of men is unavailable for no pins to be produced at all. However, in conditions where an individual can be confident in finding other men with each of the 17 other specializations (or with the ability to switch from one to another), it thus becomes advantageous to specialize in one of the steps for producing pins. This example illustrates the point that a division of labor comes with “tradeoffs”. First, for dividing labor to pay off, an individual performing all the steps must be unable to produce an outcome with the same efficiency and in the same time as a specialist. Second, in situations where the men’s interactions are limited or the number of men is too low, becoming a specialist would lead to a worse outcome than being a generalist. The idea of division of labor, like several other concepts in economics, has made its way to biological theory. Biological entities at all levels of organization exhibit division of labor, resulting in various degrees of specialization. However, in contrast to economic theory, division of labor is placed in evolutionary theory as an outcome of natural selection rather than rational decision.

One fundamental tradeoff that all biological entities face is the investment in maintenance (e.g., escaping predators, foraging, repairing damage) and reproduction (e.g., investing in gametes, finding a mate). It represents a particular kind of division, namely *reproductive* division of labor. Intriguing examples of reproductive division of labor and a high degree of cellular differentiation are multicellular organisms. A widely accepted theory even suggests that germ-soma specialization has been key for the evolutionary transition from cellular groups to multicellular individuals (Buss, 1987; Simpson, 2012). Such transitions, where multiple pre-existing entities form a new level of organization, are examples for *Major Evolutionary Transitions* (Maynard Smith & Szathmáry, 1995) or *Evolutionary Transitions in Individuality* (ETIs) (Buss, 1987; Michod, 2000). This chapter focuses on a theoretical model addressing the role of a tradeoff between life-history traits in selecting for a reproductive division of labor during the transition from unicellular to multicellular organisms (Michod, 2005; Michod, 2007; Michod et al., 2006; Michod & Herron, 2006), hereafter referred to as the life-history tradeoff model for the emergence of division of labor, or “life-history model” (*LHM*) for short.

The *LHM* has been inspired by the volvocine green algae, a taxonomic group, where today’s species range from unicellular over simple multicellular to fully differentiated (Kirk, 1998). These phototrophic eukaryotes use flagella to stay in the photic zone of freshwater environments, where photosynthesis is possible. The best-studied unicellular representative of this group is *Chlamydomonas reinhardtii*, which can be observed to possess its two flagella only for parts of its life cycle, during the growth phase. For cell division, i.e., reproduction, the flagella need to be absorbed as cells face the functional constraint of simultaneous cell division and flagellation. This constraint leads to a fundamental tradeoff between swimming and cell division (Koufopanou, 1994).

The constraint that simultaneously bears upon viability, i.e., swimming, and reproduction has been ‘solved’ in the closely related multicellular species *Volvox carteri*, where these incompatible functions are segregated into two different cell types. Its spherical colonies move around in the water column due to about 2000 cells that look very similar to the cells of *C. reinhardtii* in that they each possess two flagella. Importantly, these cells never lose their flagella and cannot divide – they are the irreversibly differentiated soma. Reproduction is carried out by a few germ cells, called gonidia, which do not possess flagella but do possess the ability to divide. In contrast to the single-celled *C. reinhardtii*, for which these two functions are temporally separated during its life cycle, the multicellular *V. carteri* displays a spatial rather than temporal separation of somatic and reproductive functions with two cell types, which is characteristic of a reproductive division of labor.

The example of the origin of the division of labor in the volvocine green algae illustrated here is not unique. In fact, the need to accommodate two incompatible processes is also thought to drive the origin of the reproductive division of labor in other multicellular groups. For example, in metazoans, the incompatibility between cell division and flagellation (King, 2004) and in cyanobacteria the incompatibility between fixation of atmospheric N_2_ and photosynthesis (Rossetti et al., 2010; Hammerschmidt et al., 2021).

The *LHM* relies on five key assumptions concerning the relationship between fitness and life-history traits (viability and fecundity) of cells and collectives: (1) Fitness is the product of viability and fecundity, (2) Collective traits are linear functions (sum or average) of their cell counterparts, (3) There is a tradeoff between a cell’s viability and its fecundity, (4) Cell traits are optimal in the sense that they display the traits that ensure the highest contribution to collective fitness, and finally (5) The viability-fecundity tradeoff is convex for large collectives due to the initial cost of reproduction. Assumptions 1-2 are summarized in Figure 1, Assumptions 3-4 in Figure 3, and Assumption 5 in Figure 4. The definitions of the symbols used are presented in Table 1; the assumptions are summarized in Table 2.

**Table 1:**
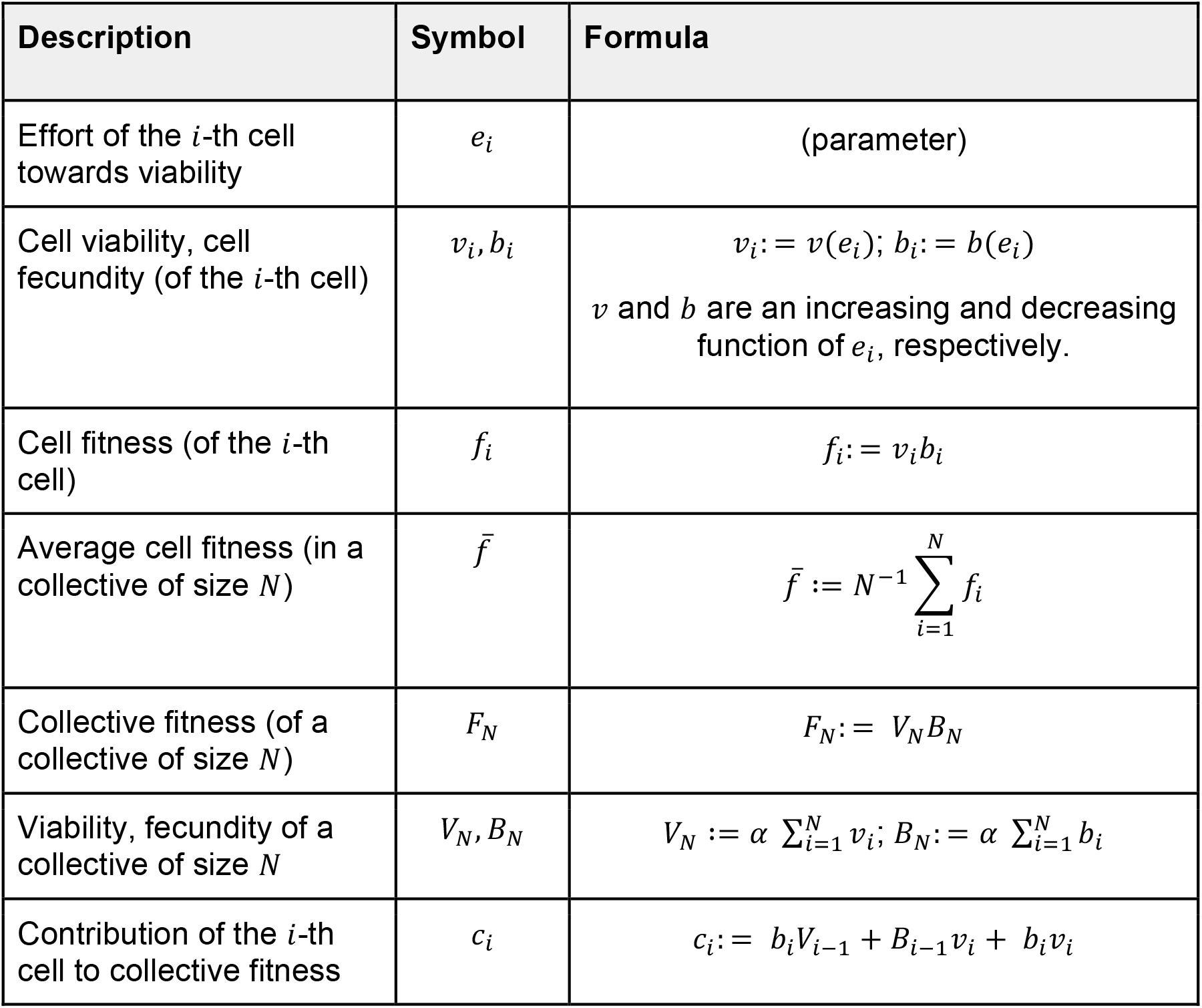
Definitions of the symbols.

| Description | Symbol | Formula |
| --- | --- | --- |
| Effort of the $i$-th cell towards viability | $e_i$ | (parameter) |
| Cell viability, cell fecundity (of the $i$-th cell) | $v_i, b_i$ | $v_i := v(e_i); b_i := b(e_i)$<br>$v$ and $b$ are an increasing and decreasing function of $e_i$, respectively. |
| Cell fitness (of the $i$-th cell) | $f_i$ | $f_i := v_i b_i$ |
| Average cell fitness (in a collective of size $N$) | $\bar{f}$ | $\bar{f} := N^{-1} \sum_{i=1}^N f_i$ |
| Collective fitness (of a collective of size $N$) | $F_N$ | $F_N := V_N B_N$ |
| Viability, fecundity of a collective of size $N$ | $V_N, B_N$ | $V_N := \alpha \sum_{i=1}^N v_i; B_N := \alpha \sum_{i=1}^N b_i$ |
| Contribution of the $i$-th cell to collective fitness | $c_i$ | $c_i := b_i V_{i-1} + B_{i-1} v_i + b_i v_i$ |

**Table 2:**
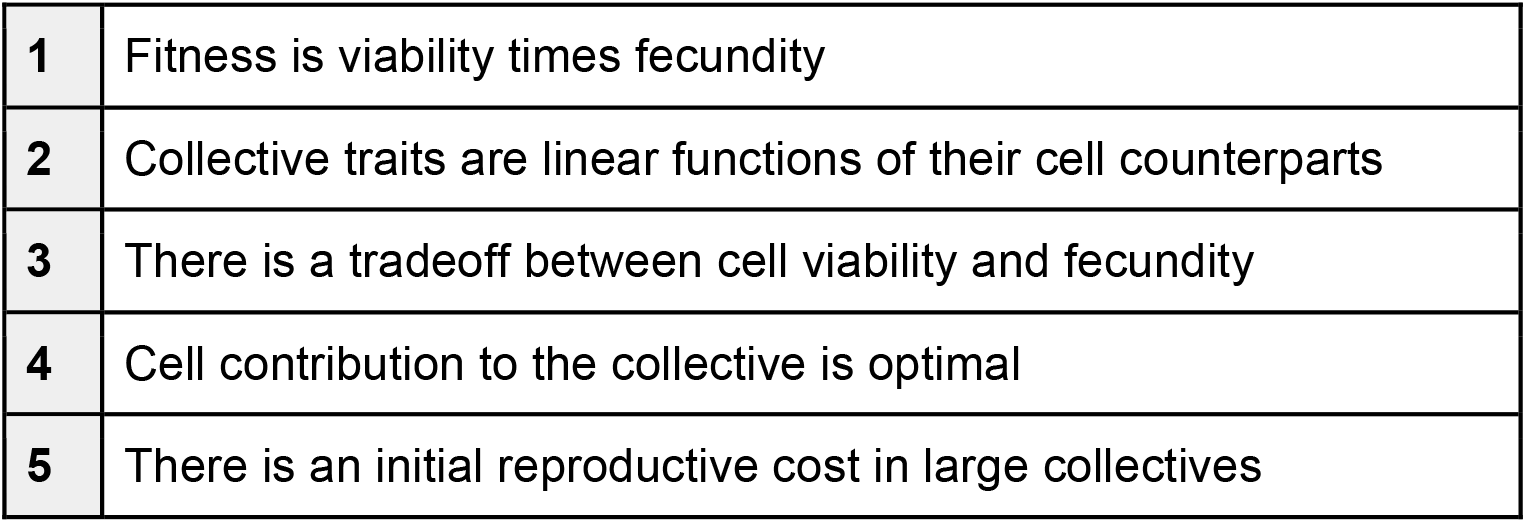
Summary of the modeling Assumptions 1-5.

**Figure 1.**
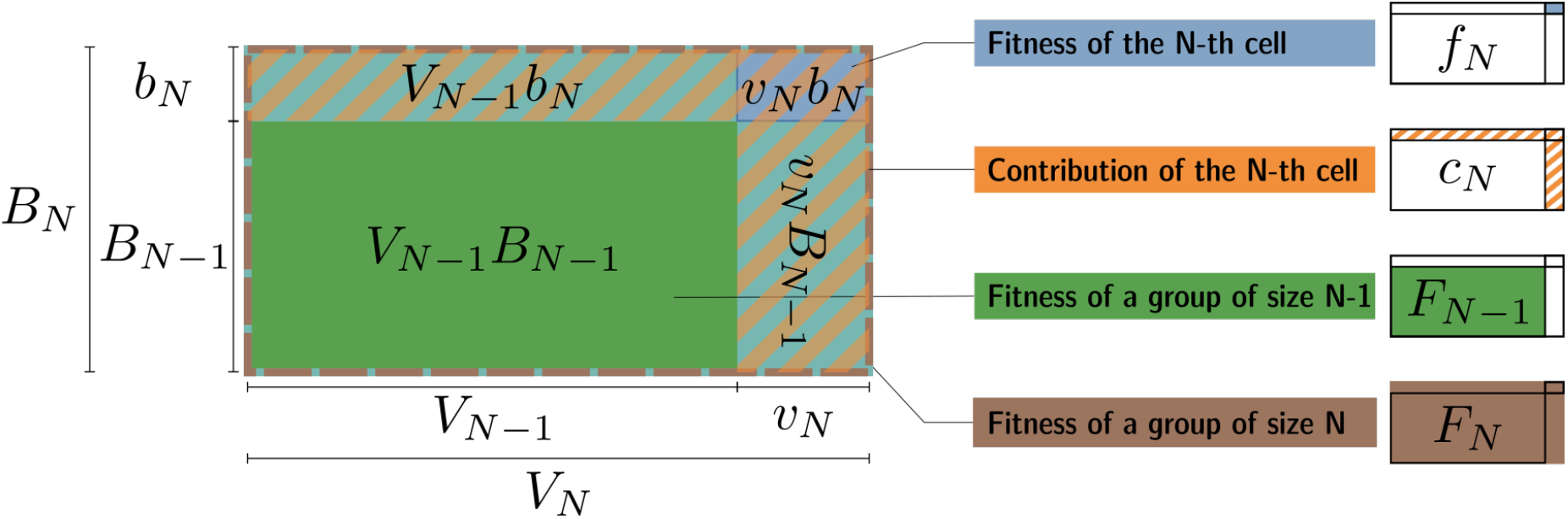
Geometric representation of collective fitness. As a consequence of Assumption 1 (fitness is the product of fecundity and viability) and Assumption 2 (collective traits are linear functions of their cell counterparts), it is possible to geometrically represent the fitness of a collective as the area a rectangle whose sides are its fecundity and viability and to decompose it into the contribution of its constituent cells. Symbols are defined in Table 1.

In this chapter, we pursue two aims. First, we provide a step-by-step guide to these assumptions for the reader to build an intuitive understanding of the *LHM*. In doing so, we highlight some of the strengths and limits of the model and provide directions to explore. Second, we present two interpretations of the *LHM* in the context of ETIs, with a particular focus on the metaphorical notion of “fitness transfer” and its limitations. Throughout, we illustrate our points with biological examples.

### 2. A step-by-step guide to the life-history model (*LHM*) of division of labor

#### 1. Fitness is viability times fecundity

Assumption 1 of the *LHM* is that the value of two life-history traits characterizes any entity (such as a cell or a collective): their viability (*v* for cells and *V* for collectives, which measures their propensity to survive) and their fecundity (*b* and *B*, which measures their propensity to reproduce). Fitness can be defined as the product of those two components (*f* = *vb* and *F* = *VB*). The effect of fitness components^1^ on the evolutionary success of organisms is at the center of life-history theory (Stearns, 1992), notably through the study of the constraints that link them together (see Stearns, 1989; and Assumption 3). Other components of fitness exist in life-history theory. However, the *LHM* focuses solely on viability and fecundity.

Taking the viability and fecundity product to compute fitness is a common assumption in the literature (Sober, 2001). There are at least two ways to justify this choice: phenomenologically and mechanistically.

First, the product of two quantities *phenomenologically* characterizes the way these two components interact in the context of fitness. One can geometrically visualize fitness as the area of a rectangle whose sides’ lengths are *v* and *b* (Figure 1). This representation helps to understand why 1) to be maximal, a multiplicative function requires a “strong balance” (Michod et al., 2006) between the two components; and 2) if one of the two sides is smaller, the marginal benefit (the surface gain) of increasing the other side is also relatively small. Additionally, if either side (fecundity or viability) has length zero, the area (fitness) is nil (we will come back to this point below).

Second, the product between two terms measuring fecundity and viability also naturally arises in various *mechanistic* models of population dynamics. As an example, consider a simple deterministic two-stage model with newborns and fertile adults that all share the same traits. Consider furthermore that whether proportion *v* of individuals that reach the reproductive stage is given by their viability *v* (0 < *v* < 1), and that all adults leave a number of offspring equal to their fecundity (*b* > 0). It follows naturally that, on average, an individual will have *vb* offspring and that the population size will grow geometrically with ratio *vb* in each generation (provided that generations are not overlapping). This growth rate *vb* is also called the *Malthusian parameter* of the population and is commonly identified as a fitness measure (Fisher, 1930, p. p22). However, in a real biological situation, there is typically no fixed proportion of the population dying at each generation, and individuals leave a varying number of offspring. Despite this, the product fecundity-viability (or equivalent ratio of fecundity to mortality) is not just a feature of simple models. It also appears in more complex, stochastic models, where those fluctuations are taken into account (Haccou et al., 2007; Kot, 2001).

In the *LHM*, any relevant entity is characterized by fitness, which is broken down into its components. In order to study the two-level system of cells and collectives of interest for the evolution of multicellularity, one has to describe how those two levels relate to each other. This is the purpose of Assumption 2.

#### 2. Collective viability and fecundity are a linear function of cell viability and fecundity

Assumption 2 of the *LHM*, which is perhaps the most controversial one, is that a collective’s viability and fecundity are considered proportional to the sum or average of its component cells’ viability and fecundity, respectively. In other words, the relationship between cell and collective fitness components is considered to be linear. Thus, a collective composed of *N* cells indexed 1,2… *N* will have the viability 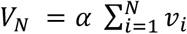 and the fecundity 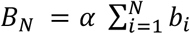 where *α* is a coefficient of proportionality. If we assume *α* = 1, it means that the collective trait is the sum of the individual traits (Michod et al., 2006). If *α* = *N*^−1^, the collective trait is the average individual trait (Michod, 2006). The value of the coefficient is a matter of simplifying expressions, and is not relevant for most results unless comparing collectives of different sizes. For ease of presentation in the rest of the chapter, let *α* = 1. The assumption of linearity is of great significance in the construction of the *LHM* because it qualifies the relationship of traits (and thus fitness components) between the lower level (cells) and the higher level (collectives). As a consequence, it allows to unambiguously define the *contribution* of the *N*-th cell to collective fitness (*c*_*N*_).

Assuming *F*_*N*_ = *F*_*N−1*_ + *c*_*N*_, we have:

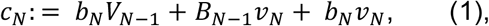

where *F*_*N−1*_, *V*_*N*−1_ and *B_N_*_−1_ respectively refer to the fitness, viability, and fecundity of a collective composed only of the cells 1 to *N* − 1. We can see that *c*_*N*_ is a sum of three terms. The first term on the right-hand side is the effect of the focal cell’s fecundity in the context of the remainder of the collective’s viability (*b*_*N*_*V*_*N*−1_). The second term is the effect of the focal cell’s viability in the context of the remainder of the collective’s fecundity (*B*_*N*−1_*V*_*N*_). Finally, the last term is the *N*-th cell’s fitness (*b*_*N*_*V*_*N*_ = *f*_*N*_). These three terms can be visualized as the sum of the three blue rectangles in Figure 1 and the contribution as the hatched orange area.

It follows from Assumption 2 that the only way a cell can affect collective viability and fecundity (and ultimately fitness) is by its own viability and fecundity. Thus, the indexing order and relative position of cells are irrelevant in this model. Furthermore, since the cells’ indexing is purely formal, any cell’s contribution to the collective can be computed in the same fashion. However, a cell’s contribution is not limited to its own fitness (the third term on the right-hand side of Equation 1). This is because it depends on the traits carried by the remainder of the collective (the first and second terms in Equation 1). Thus, the contribution of a cell can be higher if one of its components “compensates” for the weakness of the other component at the collective level (or more accurately the *N* − 1 other cells). As previously, this can be visualized in Figure 1. The same *v* quantity leads to a larger area with the orange dotted border (the cell contribution) if *B* is large and *V* is small as compared to a large *V* and a small *B*.

One consequence of Assumption 2 is that a cell with nil fitness (*vb* = 0) does not necessarily have a nil contribution towards collective fitness. For instance, consider a cell with a zero viability and fecundity of one. In this situation, only the last two terms of Equation 1 are nil. This result might look puzzling at first, especially considering that cells with nil fitness might never exist or subsist in the population (provided further implicit but standard assumptions on population dynamics) (Godfrey-Smith 2011, Bourrat 2015). However, it means that even if a cell with nil fitness (or which tends towards zero) would quickly die, its contribution to collective fitness does not necessarily tend toward zero.

A cell’s viability/fecundity contribution to collective fitness can be visualized by drawing isolines of fitness in the trait-space (see oranges lines in Figure 3). An isoline of fitness is a curve in the space *v, b* that corresponds to a fixed value of collective fitness. An isoline can be thought of as the contour lines of a map. This allows us to quickly visualize the potential contribution of any cell (i.e., any pair *v, b*) to an already existing collective. Note that the isolines are convex (see Box 1), and, provided that traits of the other cells are “balanced”, form a “hill” with its crest following along the first diagonal, when the two traits are balanced (*v* = *b*), and a valley close to the two axes, when one of the traits is close to zero. The minimum contribution is the point (0,0) where it is null, and thus *F*_*N*_ = *F*_*N*−1_.

Another way to visualize how cells with low fitness can “compensate” for each other and yield a high collective fitness is through what has been named the *group covariance effect* (Michod, 2006). Rewriting the terms of the definition of collective fitness (Table 1) shows the relationship between *F*_*N*_ and the average cell fitness,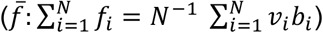 We have:

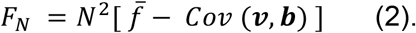

Equation 2 shows that collective fitness is not simply proportional to the average of cell fitnesses 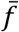,but that there is a corrective term due to the interplay of cells that can be identified as the sample covariance between the fecundity and viability of the *N* cells, which is defined as:

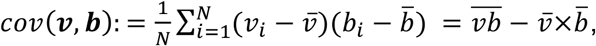

with 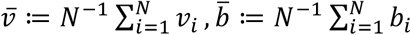 and 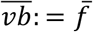 and noting that 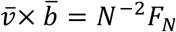

Equation 2 shows that when covariance is nil, such as when all cells are phenotypically indistinguishable (Figure 2a) or with independent trait values, collective fitness is directly proportional to the sum of its constituent cells fitnesses. However, when cell fecundity and viability are not independent of one another, covariance is not nil: it is either positive or negative. If it is positive (Figure 2b), the cells with a high *v* also have a high *b*, resulting in what we call “all-or-nothing cells”. The opposite is true if it is negative (Figure 2c), resulting in specialized germ or soma-like cells.

**Figure 2.**
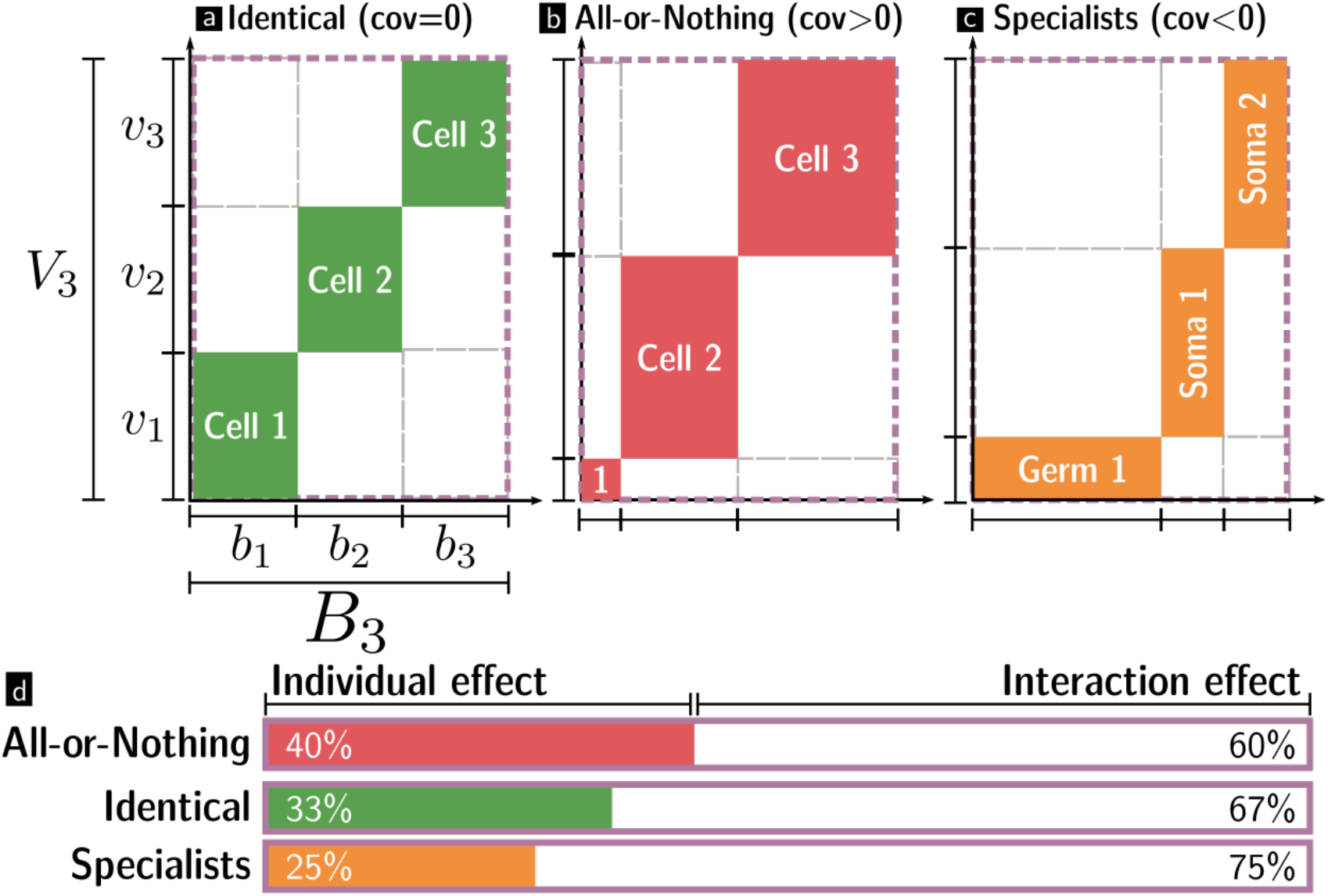
The covariance effect quantifies the extent to which collective fitness depends on intrinsic and interaction effects between cells. Three collectives composed of three cells are represented in the viability-fecundity space. The traits of each cell *v*, *b* are represented in color, and their fitness *f* is the area of a colored rectangle (green, red, orange). The three collectives have the same fitness *F*_3_ = *V*_3_*B*_3_, represented by the area of the purple rectangle and the same average cell fitness 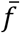. The collective fitness *F*_3_ is the sum of the cell fitnesses (colored tiles, *f*) and of the interaction effects between cells (in white). **a)** A collective composed of three identical cells with equal traits (null covariance between v and b). **b)** A collective composed of “all-or-nothing cells” that would simultaneously have a high (low) viability and fecundity (positive covariance) (these cells are conceptual constructs; they are not biologically plausible). **c)** A collective composed of specialist cells with high fecundity and low viability (germ) or low fecundity and high viability (soma). **d)** The sum of cell fitnesses (colored area, labeled “individual effect”) represents a larger fraction of the collective fitness (purple delimited area) when cells do not compensate for each other’s weaknesses (all-or-nothing cells) as compared to when they do (specialists).

Cell-cell interactions compensate for cell heterogeneities only when the covariance is negative. This can be seen by tallying the relative weight of “individual effects” of the cells on collective fitness (i.e., the direct sum of cell fitnesses, in color in Figure 2) and “interaction effects” due to the cross product between cell traits (the rest, in white in Figure 2). When done, it becomes apparent that individual effects are relatively more important when the covariance is positive and relatively less important when the covariance is negative (Figure 2d). The converse is true for the interaction effects. Interaction effects are important because they explain how a collective can have high fitness, even if the fitness of its constituent cells is constrained to be low.

As stated earlier, Assumption 2 characterizes the relationship between cell and collective fitness in a more subtle way than just taking the average. It also permits to study the combined effects of any set of cells (characterized by viability-fecundity pairs), as well as to tease apart a cell’s direct contribution and its interactions with other cells in the collective by using the cell contributions (Equation 1) and the covariance effect (Equation 2). However, Assumption 2 is quite strong and thus comes at a steep price. In particular, it limits the range of phenomena that can be satisfyingly described by the model. As discussed below, biologically plausible scenarios of non-linear and non-monotonic or in general higher-order interactions are impossible to describe within this framework because of this assumption. This limitation should be kept in mind by experimentalists and modelers alike.

A further limitation of Assumption 2 comes from the fact that it implies a monotonic relationship between cell and collective fitness. Thus, increasing the fecundity of one or all the cells of a collective of a given size is assumed to *always* increase the whole collective’s fecundity by the same magnitude (up to the proportionality coefficient). In turn, this brings a net collective fitness increase, even though the return might be diminishing (when viability and fecundity are not well balanced). We can imagine that this assumption might not hold for all trait values. Increasing cell fecundity might increase collective fecundity by increasing the potential number of propagules this collective can produce. However, we might reasonably think that the fast proliferation of cells negatively interacts with the propagule-producing mechanisms when above a certain threshold.

Related to the previous point, Assumption 2 also implies a kind of “beanbag” model of collectives, where the relative position and orders of cells cannot be captured. It might seem obvious for eukaryotes with sophisticated developmental dynamics and organ partitioning that a cell will have a different impact on the collective fate depending on its position and the nature of its neighboring cells. However, even relatively simple examples of multicellular organisms, such as heterocyst-forming filamentous cyanobacteria, show why this is pervasive. In these species, the lack of combined nitrogen in the environment induces the formation of differentiated cells, heterocysts, which are devoted to the fixation of atmospheric N_2_. Heterocysts exchange fixed nitrogen compounds against carbon products with the neighboring photosynthesizing (vegetative) cells of the filament. Importantly, heterocysts are not located on random spots in the filament but are spaced at regular intervals (Yoon & Golden, 1998). For example, in the model species *Anabaena sp*. PCC 7120, heterocysts are separated by 10-15 vegetative cells (Herrero et al., 2016). This ensures an adequate supply with fixed nitrogen compounds while maximizing the number of vegetative cells within a filament (Rossetti et al., 2010). Notably, while vegetative cells can divide and generate all other specialized cell types, heterocysts cannot divide and are terminally differentiated. Thus we do not only observe a metabolic division of labor but a reproductive one, where heterocysts are comparable to the somatic and the vegetative cells to the germ cells in multicellular eukaryotic organisms (Rossetti et al., 2010). This structure cannot be described accurately in the original *LHM* (but see Yanni et al., 2020)

Assumption 2 implies that increasing any individual trait is bound to increase its collective counterpart. Assumption 3 prevents the simultaneous increase of both viability and fecundity.

#### 3. Tradeoff between cell viability and fecundity

Assumption 3 of the *LHM* posits that a cell with a particular value for viability is necessarily constrained on its counterpart value for fecundity. Consequently, this reduces the number of free dimensions in the model: the two traits cannot vary independently.

This assumption covers the intuitive point that a cell cannot simultaneously be highly fecund and highly viable (we call “all-or-nothing cells”) if it has a finite amount of energy to allow both of these (biological) functions. There are many ways to implement a tradeoff in a model. In the *LHM*, this is done using a relatively simple, deterministic, and one-dimensional method. Consider that, besides viability and fecundity, there is a third “hidden” trait for a cell, noted *e* that quantifies the *effort* or *investment* towards one of the two traits. Then, by definition, the viability is an increasing function of the effort, *v*_*i*_ = *v*(*e*_*i*_), and the fecundity a decreasing function of the effort: *b*_*i*_ = *b*(*e*_*i*_). Here, *v* and *b* are (mathematical) functions that must be specified by the modeler. For instance, a simple linear tradeoff can be defined as *v*(*e*) = *e* and *b*(*e*) = 1 − *e* for *e* ∈ [0,1].

If the notion of effort is important for understanding the logic of the tradeoff, it can be abstracted graphically when representing the tradeoff in the (v,b) plane introduced in Assumption 2. The tradeoff can be represented as a curve (purple in Figure 3) constituting of all the combinations of *v*_*i*_ *b*_*i*_ given by all possible values of *e*_*i*_. Different functional forms result in different tradeoff shapes. Its shape, and in particular its convexity, are at the base of many strategies within the framework of life-history theory (See Box 1 for a primer). We will return to it when discussing Assumption 5.

The notion of tradeoffs in life-history theory is an indubitably elegant way to incorporate an organism design’s underlying constraints into a model. For instance, it can be used to take into account that the microtubules organizing center in the *Volvocaceae* cannot simultaneously take part in reproduction (through mitosis) and viability (through flagellar motility) (Koufopanou, 1994). While they are powerful theoretical tools, the existence of tradeoffs is hard to demonstrate, let alone to quantify. One reason is that they can originate from many sources: physical (diffusion, buoyancy), genetic (metabolic pathways, regulations), or ecological (grazing, parasites) constraints. Moreover, tradeoffs are not always set in stone. If physical constraints like diffusion hardly change, mutation events can overturn other constraints: for instance, in the flagellate *Barbulanympha*, the microtubule organizing center can simultaneously participate in locomotion and reproduction (Buss, 1987). Note that in the *LHM*, the shape of the tradeoff changes with the size of the collective. This will be treated in more detail as part of Assumption 5.

Following Assumption 3 the set of all possible cells is reduced, as the trait of any new cell must be located on the tradeoff curve. The model is not complete yet; natural selection acts on the organism in the context of these tradeoffs, and its effect must be described. This is the purpose of Assumption 4.

#### 4. Cell contribution to the collective is optimal

So far, the role of natural selection has seldom been invoked in the *LHM*. We have only described the property of cells and collectives and the diversity of traits they can exhibit, given some underlying constraints. Assumption 4 models the consequence of natural selection on this system: it assumes that all cells are optimal in terms of their contribution *c* to collective fitness. Formally, it means that the life-history traits of any cell *i* within the collectives are such that the value of *c*_*i*_ is maximal:

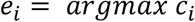

Note that optimality is an *assumption* rather than an outcome of the model.

Graphically, to find the values for a cell to contribute optimally to the collective, one has to look for the intersection between the tradeoff curve (purple in Figure 3) and the highest isoline of collective fitness (orange in Figure 3). This point is where a cell existing within the physiological constraints that link *v* and *b* has the highest fitness contribution. If the shape of the tradeoff is simple enough, there is a single optimal point, and thus the model predicts the traits of any new cell-based on Assumptions 1-4 plus the shape of the tradeoff.

**Figure 3.**
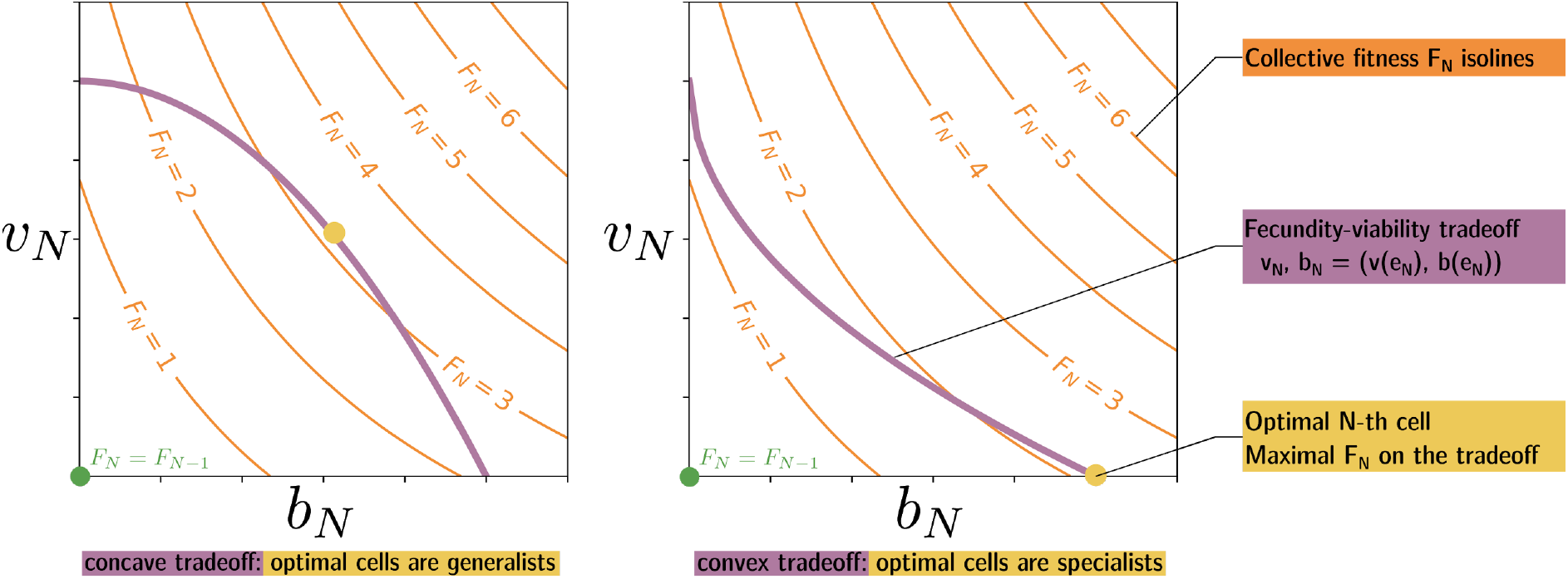
Isolines of fitness, tradeoffs, and optimality of cell contributions. Representation in the plane (*b*_*N*_, *v*_*N*_) formed by the fecundity and viability of the N-th cell of a collective. As a consequence of Assumptions 1 and 2, collective fitness is a surface (represented by orange isolines) and, at the origin (*v*_*N*_ = *b*_*N*_ = 0), the collective fitness is minimal (*F*_*N*_ = *F*_*N*−1_, green dot). As a consequence of Assumption 3, the values of *v*_*N*_ and *b*_*N*_ is constrained by a tradeoff (purple line). As a consequence of Assumption 4, the model predicts that the traits favored by natural selection are the ones that yield the highest collective fitness (yellow disk) while satisfying the tradeoff constraint. Concave (and linear) tradeoffs (left) favor generalists cells (with balanced *v*, *b*), while convex tradeoffs (right) favor specialist cells (with high *v* and low *b*, or vice-versa).

Of course, Assumption 4 could turn out to be incorrect if the cells are *not* optimal in their contribution to collective fitness. Cells might not be optimal for several reasons. For instance, the optimization of their traits might not occur independently from one another because they share the same underlying developmental programme. Alternatively, they might be “stuck” in another region of the trait space (no viable mutation path would bring them to the optimal phenotype). Other reasons include that the tradeoff’s shape has recently changed due to changes in the environment or that evolutionary forces (such as selection at another level or an evolutionary branching point) prevent the cells from reaching or remaining at the optimal phenotype.

Thus far, we have seen that the *LHM* assumes that the reproductive success of a collective depends on two fitness components (Assumption 1) that derive from their cell counterparts (Assumption 2), which are linked by underlying constraints (Assumption 3), and that natural selection is expected to favor optimal cells within this context (Assumption 4). The last piece of the puzzle is to qualify the shape of the tradeoff. Assumption 5 does precisely this.

#### 5. There is an initial reproductive cost in large collectives

Assumption 5 states that small collectives have a linear or concave tradeoff—favoring generalist cells—while large collectives have a convex tradeoff—favoring division of labor. The distinction between linear, concave, and convex tradeoffs is presented in Box 1. The mechanism proposed to explain why large collectives have a convex tradeoff is the initial cost of reproduction. This assumption is important because it characterizes the underlying constraints that bear on cell traits, but at the same time, tie them to the collective, in particular to collective size.

To understand Assumption 5, consider a cell specialized in viability (i.e., with a low fecundity) (Figure 4a). The mechanism for the initial cost of reproduction hinges on the assumption that if this cell was investing more in fecundity than it currently does, it would reduce its viability but would *not* increase its fecundity (Figure 4b) until a threshold is reached (Figure 4c) after which it would increase (Figure 4d) until the cell is fully specialized in fecundity (Figure 4e).

**Figure 4.**
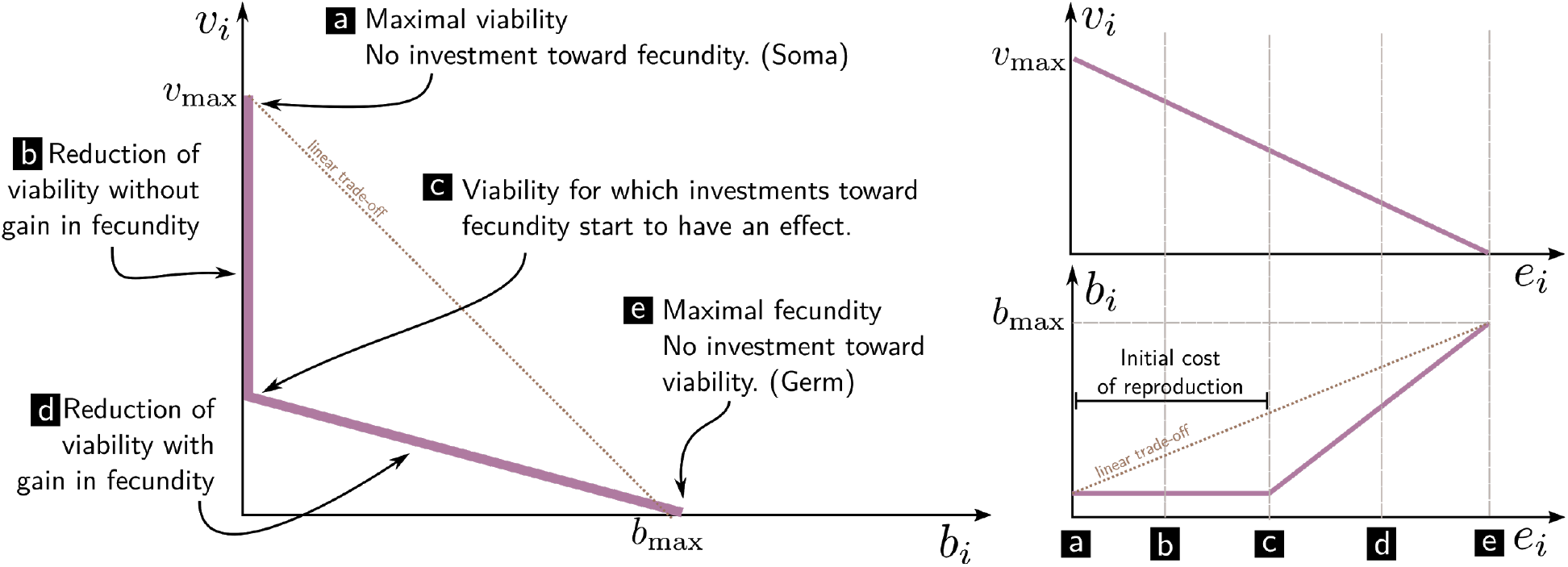
Initial reproductive cost as a model for a convex tradeoff. Cell viability *v*_*i*_ and fecundity *b*_*i*_ are constrained, as represented by the purple curve corresponding to the possible combinations (*v*_*i*_, *b*_*i*_) under the tradeoff modeled by the reproductive effort *e*_*i*_. Starting from maximal investment towards viability (*v*_*max*_, a), reduction in the viability effort as no effect on fecundity (b) until a threshold (c), where fecundity increases (d) up to the maximal fecundity allowed by the model (*b*_*max,*_ e). Contrast this with the simple linear tradeoff (brown).

The relationship between group size and the shape of the tradeoff between contribution to collective viability and fecundity is generally understood in terms of physical constraints. For instance, at the collective level, in the volvocine green algae, the enlargement of reproductive cells increases the downward gravitational force, increasing sinking and is only overcome by the investment in more buoyant somatic cells (Solari et al., 2015). Thus, when colony size increases, a required initial investment toward buoyancy appears that did not exist in single-cell organisms. This, in turn, explains how the tradeoff, taken to be linear (or even concave) for single cells and small collectives, becomes convex when considering larger groups.

This is the last part of the *LHM*. As a consequence of Assumptions 1-5, large collectives favor the selection of specialist cells, and thus division of labor.

**Box 1. Tradeoff convexity**

A tradeoff is a relationship linking two quantities that cannot simultaneously be maximal: often, if one increases, the other must decrease. In this chapter, those two quantities are the life-history traits of an individual (viability *v*, and fecundity *b*).

This can be due to a variety of phenomena: a tradeoff between size and nutrient intake might result from physical laws, such as diffusion, or the tradeoff may come from the resource allocation of an organism (with a given quantity of nutrient, only so many molecules might be synthesized, creating a natural tradeoff between structural molecules, housekeeping, and reproductive machinery), tradeoffs may also be caused by the underlying genetic structure of the organism (e.g., a single regulator molecule acting on two pathways, making regulation of one and the other correlated, or the functional constraint of simultaneous cell division and flagellation in the case of *C. reinhardtii*), or through interaction with other species (the expression of a useful transporter might render the cell vulnerable to a certain type of virus). As a consequence, tradeoffs themselves might change during the evolutionary history of organisms.

This box gives a short introduction to the simple, deterministic one-dimensional tradeoffs that are used in the *LHM* for the division of labor in multicellularity. Additional resources can be found in life-history theory textbooks, such as Flatt (2012). Two-dimensional tradeoffs are conveniently represented by putting the two measures on the axis of a plane and shading the area of pairs of values that are possible within the confine of the tradeoff (Figure B1). This may result in a surface (two degrees of freedom) or a curve (one degree of freedom) depending on the number of free dimensions the tradeoff allows.

Such a tradeoff provides a straightforward definition of a “specialist” organism (with a maximal or close to the maximal value in a trait and accordingly a lower value for the other trait) and a “generalist” organism (with an intermediate value in both traits).

**Figure B1:**
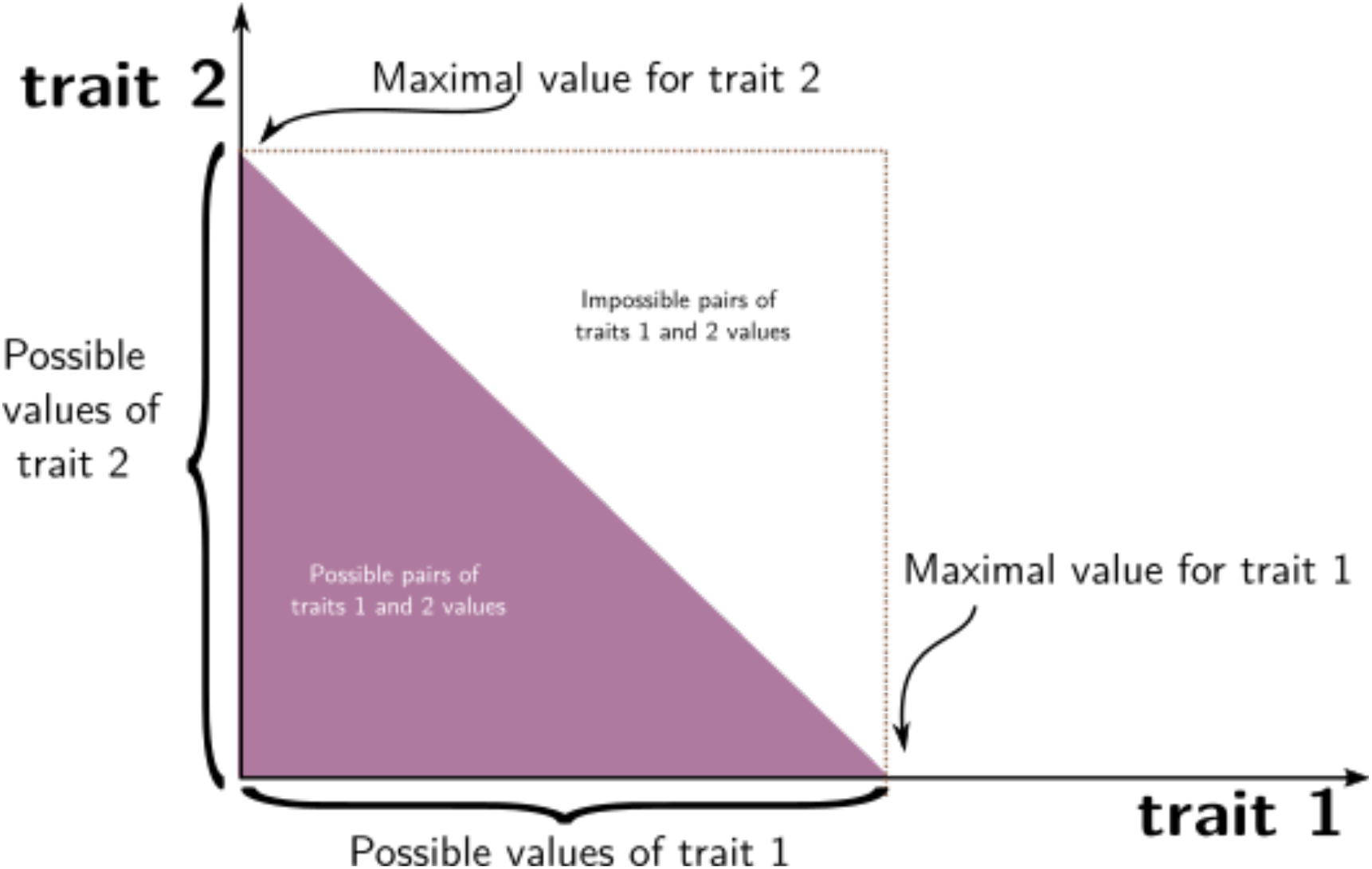
Example of a tradeoff between trait 1 and trait 2 with 2 degrees of freedom, represented by the purple surface.

A particularly important graphical way of analyzing a tradeoff is to consider its position with respect to any segment defined by any two couple of trait values. The tradeoff curve (or the edge of the surface) might coincide (Figure B2.a), go below (Figure B2.b), above (Figure B2.c), or cross (Figure B2.d) these segments.

When the curve coincides with all segments, the tradeoff is said to be linear. In this case, the relation between the two traits is proportional, reducing the value of trait 1 by a quantity *x*, and results in increasing the value of trait 2 by *ax* (where *a* depends on the slope of the curve and may depend on the scaling of the trait values). When the curve is below all segments, the tradeoff is said to be convex: A small reduction in trait 1 has a different effect on trait 2 if the trait is close to the maximum value (small effect when compared to the linear) or to the minimum (large effect). Similarly, if the tradeoff is above all segments, it is said to be concave. In this case, a small reduction in an optimal trait has a large effect on the other trait, whereas a small increase in a low trait has a small effect on the other trait. If the curve is above some segments and is below or crosses others, it is neither convex nor concave but can be studied in part by focusing on the different regions.

**Figure B2:**
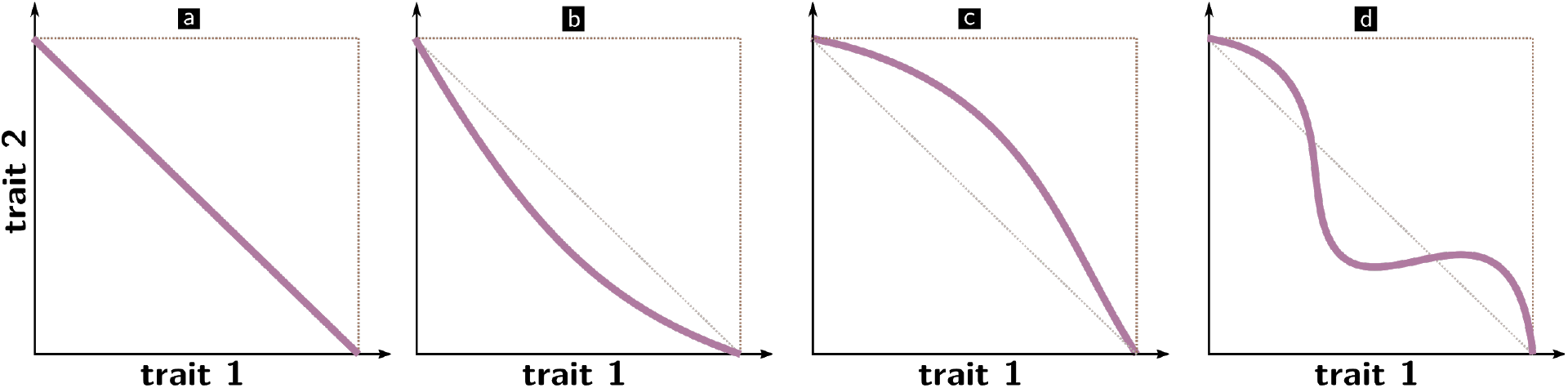
a) Linear, b) convex, c) concave, d) composed tradeoff.

Intuitively, if the tradeoff is convex, being a specialist (i.e., being on either axis) is the only way to reach high trait values, while being a generalist is “costly” in the sense that it comes with a large reduction in trait value. This is opposed if the tradeoff is concave: generalists enjoy a less pronounced reduction of their trait values with respect to specialists (being a specialist can be considered “costly” in the sense that the marginal cost of increasing a high-value trait is relatively high compared to the case of a convex tradeoff).

Note, however, that the convexity (or even the shape) of a tradeoff does not make a prediction about the outcome of the evolutionary process on its own. It just delimits the set of possible organisms. To be able to predict the outcome of the evolutionary process from such a tradeoff, one has to make additional assumptions. For instance, one could assume, as we do in the *LHM*, that a fitness function F of the two traits exists and that the evolutionary dynamics reached an equilibrium state in which only the organisms with the highest fitness F are represented in the population. (In this case, one needs to look for the value within the set of possible individuals that gives the highest fitness). If density-dependent interactions are suspected of playing a role, a possibility may be to define the invasion fitness of a rare mutant in a resident population for all pairs of points in the tradeoff and look for evolutionary stable strategies (ESS), following the adaptive dynamics method (Geritz et al., 1998).

### Discussion: Fitness interpretations in ETIs

The previous section presented a mechanism that promotes cell specialization between two life-history traits (viability and fecundity), hence a division of labor. This mechanism is based on the presence of a convex tradeoff between the two traits due to the existence of an initial cost to reproduction in large collectives. This section puts this model back in the broader context of ETIs by contrasting two interpretations of this phenomenon.

### ‘Reorganization and transfer of fitness’ interpretation

A first interpretation of the *LHM* is based on the idea that the hallmark of an ETI is fitness reorganization/transfer/decoupling in the sense that cell specialization results in the lower-level cells “relinquishing their autonomy in favor of the group” (Michod, 2005; Michod et al., 2006), resulting in a transfer of “fitness and individuality [from] the cell level to the group level” (ibid). This is the interpretation we have tacitly assumed throughout because it the one with which the *LHM* was initially proposed when interpreting *f*_*i*_ as cell fitness and *F*_*i*_ as collective fitness.

This interpretation is rooted in the Multi-Level Selection 1-2 framework (Damuth & Heisler, 1988; Okasha, 2006) as cited in (Michod, 2005). In Multi-Level Selection 1 (MLS1) models, collective fitness is taken to be the average fitness of its members (or proportional to it), while in Multi-Level Selection 2 (MLS2), collective fitness cannot be defined in terms of particle survival and reproduction. As a result, a new notion of fitness must be devised.

From this interpretation, the problem of explaining ETIs can be “reduced” to explaining the transition from an MLS1-like situation to an MLS2-like situation (Okasha, 2006, Chapter 8). This may initially seem like an insurmountable hurdle because in MLS1 (before the transition), cells are selected to have the highest cell fitness. In contrast, fully specialized cells in the model (after the ETI has occurred) have nil (or close to nil) fitness. The concept of *fitness transfer* (Michod, 2006) solves this problem by considering that during an ETI, fitness between the two levels is reorganized: it is transferred from the lower level (the cells) to the higher level (the collective). The transfer is achieved through germ and soma specialization (which results from the tradeoff’s convexity). When cells become specialized, they relinquish their fitness to the benefit of the collective. Furthermore, during the process, collective fitness transitions from a mere average of cell fitness (MLS1) to a quantity, which is no longer the cell fitness average in the collective (MLS2) because of the covariance effect.

While this interpretation is appealing, there are some problems associated with it. The main reason is that it seems to imply that fitness is a material quantity that can be transferred from one entity to another, comparable to a liquid that can be poured from one container to another. While, at first glance, this analogy may seem a helpful image to get an intuitive idea of the problem, it contradicts our modern understanding of fitness as a predictor of evolutionary success (Bourrat, 2015a, 2015b; Doulcier et al., 2021). First, it implies that some cells with nil fitness or close to it are not dead, contradicting the principle of natural selection. Second, because the evolutionary fates of cells and collectives are tied (by virtue of being made of the same biological substrate), it is far from obvious to see how one level could ever be favored at the other’s expense (Black et al., 2020; Bourrat, in press). This point is known in the philosophical literature as the “causal-exclusion principle.” If a phenomenon is explained or described exhaustively at the lower level, one cannot appeal to the higher level to further explain this phenomenon. Doing so is either a form of “double counting” or requires assuming that strongly emergent properties are created ex-nihilo at the higher level. Assuming the existence of strongly emergent properties raises a new range of issues because they contradict with materialism, the idea that all causes are physical in nature.

Finally, the fitness transfer interpretation implies that cells constituting a multicellular organism can have different fitness values. This conflicts with the fact that those cells are clones and should thus have the same (inclusive) fitness (Bourrat, 2015b). To make this point clear, while a queen and a worker bee have different reproductive outputs, they have the same (inclusive) fitness. Evolutionarily, it does not make sense to say that the queen is more successful than the worker. Similarly, it does not make sense to say that a liver cell in a multicellular organism is fitter than a brain cell.

Recent work has helped solving these issues by proposing a new interpretation of fitness at different levels of organization (Shelton & Michod, 2014, 2020). Following this new interpretation, the term “cell fitness” does not represent the cell’s fitness within the collective but rather the one it *would have* if it were without a collective (counterfactual fitness). When cells have the same fitness they would have in the absence of collective, no transition has taken place. However, when the cells have a different fitness, a transition has occurred (or at least has been initiated). This constitutes a reasonable argument toward deciding whether an ETI happened (i.e., the state of cells and collectives). Crucially, this says nothing about the mechanism of the transition. There is no actual “decoupling” or “transfer” of fitness, other than in the loose metaphorical sense that the purely theoretical counterfactual fitness aligns or not with the actual fitness.

Metaphors and analogies are incredibly useful in biology because they allow us to build intuition of complex mechanisms by drawing parallels with other systems. *Fitness transfer* or *reorganization* implies the physical transfer of a material quantity (with or without conservation). However, fitness is not transferred from one place (the cell) to another (the multicellular organism) in a way heat can be transferred. “Fitness transfer” might be used, but only in a loose metaphoric sense, that is, in the same way teleological language in evolutionary biology is used in the context of a teleonomic explanation (Jacob, 1970; Pittendrigh, 1958). A further point worth mentioning is that metaphors might favor a specific interpretation that could obscure some aspects of the phenomena studied, similarly as the sole focus on selection created the blindspots of the adaptationist programme (Gould et al., 1979).

### ‘Propensity’ interpretation

We favor an alternative interpretation of the *LHM*. This interpretation starts with the same mechanism: a convex tradeoff between contribution to collective fecundity and collective viability will promote the emergence of individual specialist cells, thus a division of labor. It diverges from the previous one by its treatment of fitness, emphasizing on how it emerges from cell *traits* rather than using it as a reified quantity of cells and collectives.

For the concept of fitness to qualify as a predictor in evolutionary biology, it cannot be reduced to an entity’s actual success (i.e., its realized fitness). Instead, it must be tied to its *potential* success (or success in the long run). Without this point acknowledged, fitness is condemned to be tautological, as philosophers and biologists alike have long recognized (Manser, 1965; Popper, 1974; Smart, 2014; reviewed in Doulcier et al., 2021).^2^ This conundrum led to establishing several frameworks for the interpretation of fitness, one among which is the propensity interpretation of fitness (Beatty, 1984; Brandon, 1978; Pence & Ramsey, 2013). According to the propensity interpretation, fitness is a probabilistic property of entities summarizing their probability distribution of reproductive success (as defined by their demographic parameters: birth and death rates) in a given environment.

If we adopt this interpretation, the problems raised with the fitness transfer interpretation vanish. First, the problem of a collective’s different (clonal) cells having different fitnesses disappears. Even though their *realized* fitness (actual life-history) might be different, their “true” fitnesses (potential life-history) are equal because they relate to the potential success of the same genotype. Second, this interpretation does not appeal to fitness transfer or decoupling since cell and collective fitness are computed in expectation. Following the transfer of fitness interpretation, although this is not made explicit in the model, cell fitness and collective fitness are computed relative to different environments. In particular, collective-level demography (i.e., birth and death events of collectives) is typically ignored when computing cell fitness. As a consequence, it becomes possible to define different values of fitness for the collective (*F*) and the cells (*f*). However, the fact that they are computed in different environments implies that they cannot legitimately be directly compared. When cell and collective fitnesses are computed in the same environment following the propensity interpretation, for instance, by factoring in collective events in the cell-level computation, they are necessarily equal (Bourrat, 2015a, 2015b; Bourrat et al., 2020). Cells and multicellular organisms are two levels of description of the same physical reality and cannot contradict each other despite some claims to the contrary (e.g., Okasha, 2006).^3^ Even though conflicting processes might exist (segregation distortion locus, cancerous growth), fitness, properly computed to be comparable, must tally these conflicts and be coherent when refereeing to the same entity, regardless of the method of description.

Alternatively, one way to connect the interpretation we favor and the counterfactual fitness approach is to compare the fitness of free-living cells with cells within the collectives and observe *apparent decoupling* between these two environments: a decrease in free-living fitness and an increase in within-collective fitness (Bourrat, 2015a, 2015b, 2016; Bourrat et al., 2020; Bourrat, in press). However, this apparent decoupling is rather a sign of *linkage* between the traits that contribute to free-living and within-collective fitness. The propensity interpretation of fitness can explain the same phenomena without invoking any “fitness transfer.” If there is a “transfer”, it is between the energetic investment of the cell towards different traits: from traits that provide no advantage to cells living in a collective (and potentially contributing to a free-living life-cycle), towards traits that give an advantage to the cells living in a collective (including vicarious advantages of cells with the same genotype).

Doing away with the reifying idea that fitness is something to be transferred, and more generally treating the MLS1/MLS2 distinction as conventional—i.e., two different ways to formalize the same thing—rather than as an evolutionary mechanism, allows pursuing lines of inquiries that were harder to conceive within this framework. For instance, the focus on the relationship between cell and collective fitness naturally leads to the assumption that contributions to the collective fitness component are linear functions of their free-living counterparts (as was the case in Assumption 2). However, designing a mechanistic model naturally leads to relaxing this assumption. Traits of collectives are most certainly more complex than the arithmetic aggregation of individual quantities measured in the propagule or the fully developed collective. They are rather the result of internal developmental dynamics, i.e., within-collective cellular ecological dynamics (Hammerschmidt et al., 2014; Rose et al., 2020). Selection of developmental mechanisms has long been recognized as an important part of ETIs (Buss, 1987; Michod & Roze, 1997) and can fully be studied by models that explicitly describe the ecological dynamics within collectives (see Ikegami & Hashimoto, 2002; Williams & Lenton, 2007; Xie et al., 2019 for general cell communities; but see Doulcier et al., 2020 for an application to ETIs).

## Conclusion

Division of labor is observed in complex organisms. The functions exhibited by multicellular organisms cannot be exhibited simultaneously by a single cell. Multicellularity solves this problem by allowing different subsets of cells to perform the different functions at once. The level of division of labor exhibited by a collective varies with the extent to which cells are specialized.

Natural selection favors specialist cells (hence division of labor) if there is a convex tradeoff between two equally important functions for cell fitness. The convexity of the tradeoff is a consequence of two hypotheses: first, an energetic investment model in which a cell has limited energy to invest in two traits that contribute towards each function, and second, an initial investment cost, in which a small investment in a trait does not immediately translate in an improvement of the function. The *LHM* predicts that collectives constituted of cells investing less energy in traits that contribute toward free-living fecundity and viability but more in traits that contribute towards fecundity and viability of collectives will progressively outcompete other collectives and become widespread.

This phenomenon has been interpreted as a “transfer of fitness” in the sense that individual cells relinquish their autonomy (investing less in free-living traits) to participate in life-history traits of collectives (investing more in contribution towards collective function). During this “reorganization of fitness”, cell fitness has been proposed to decrease while collective fitness increases. The fact that cell fitness and collective fitness do not change in the same direction has been named “fitness decoupling”. However, this interpretation can be misleading because it is in conflict with the concept of fitness as used in evolutionary biology. To fully appreciate the relevance of the *LHM* to ETIs, two things must be stressed. First, tradeoffs occur between traits, not between fitnesses at different levels of organization. Second, fitness can only be defined with respect to a given entity (cell or collective) in a given environment and cannot be incoherent between the whole and the part. Thus, fitness cannot literally be “transferred” from individuals to collectives, even if, in retrospect, the traits that are adaptive in a collective-environment would be detrimental to a free-living organism.

Tradeoffs between life-history traits are valid mechanisms - independently of the interpretation in terms of fitness transfer or steady-state propensity, or even any other kind of interpretation one might propose (e.g., inclusive fitness, game theory, altruism). The interpretation chosen only represents a useful narrative for placing ETIs in the broader context of the evolution of complexity and allowing us to pursue subsequent questions, e.g., regarding developmental programs. Nonetheless, invoking a fitness concept that is consistent with the broader use of this term represents the primary reason for preferring one interpretation of the *LHM* to the other.

## Acknowledgments

The authors are thankful to the Theory and Method in Biosciences group at the University of Sydney. GD’s and PB’s research was supported by a Macquarie University Research Fellowship and a Large Grant from the John Templeton Foundation (Grant ID 60811). KH is grateful for support by *The Hamburg Institute for Advanced Study* (HIAS) and the Joachim Herz Foundation.

## Footnotes

1 or life-history traits, the two terms are often used indistinguishably (Flatt, 2012).

2 The propensity interpretation of probability is contentious in philosophy (Hájek, 2012). It comes with a number of problems, some of which are inherited by the propensity interpretation of fitness (Bourrat, 2017; Godfrey-Smith, 2009). In recent years, several alternative interpretations of probabilities that play the same role as propensities and solve the issues of the propensity account have been proposed (e.g., Abrams, 2012; Lyon, 2011; Rosenthal, 2010; Strevens, 2011). Addressing the differences between these various interpretations in the context of fitness is beyond the scope of the present work. For our purpose, we use “propensity” loosely as an entity’s dispositional property to produce offspring (or equivalent terms in the aforementioned interpretations) without committing to any particular probability interpretation.

3 This claim admits a few theoretical exceptions, which are not relevant to ETIs.

